# Identification of a novel structural motif and overexpression of key transcripts elucidated in Adenovirus 10

**DOI:** 10.1101/2025.09.20.677498

**Authors:** Rosie M. Mundy, Kasim Waraich, Emily A. Bates, Pierre J. Rizkallah, Alexander T. Baker, Mark T. Young, Edward Morris, Paula C. A. da Fonseca, Carly M. Bliss, David Matthews, David Bhella, Alan L. Parker

## Abstract

Adenoviruses are widely used as vectors for subunit vaccines and oncolytic therapies. Efficient vectors must infect target cells and deliver therapeutic transgenes at high levels. Species D adenoviruses, such as human adenovirus type 10 (HAdV-D10), are promising candidates due to low seroprevalence in humans. Here, we present the cryo-electron microscopy structure of the HAdV-D10 capsid alongside transcriptomic profiling of infected cells to inform vector development. The fiber shaft, essential for cell entry, was resolved at 4.6 Å, revealing a previously uncharacterized ‘umbrella’ motif. Viral transcript analysis using an ORF-centric pipeline uncovered key differences from HAdV-C5, including abundant expression of a transcript encoding a protein equivalent to mature protein VII. These findings highlight the importance of detailed vector characterization prior to clinical translation and support the advancement of HAdV-D10 as a next-generation platform for gene delivery and vaccine development.

## Introduction

Human adenoviruses (HAdV) are double stranded DNA viruses with non-enveloped icosahedral capsids (*1*). HAdV are split across species A-G based on serology, sequence identity and pathogenicity (*2*). Species D (HAdV-D) is phylogenetically the largest species of HAdV as a result of recombination events giving rise to new adenoviruses serotypes (*3*). HAdV-D viruses often cause minor illness, including ocular infections (*1*). Adenovirus 10 (HAdV-D10) was isolated from the eyes of conjunctivitis patients with clinical symptoms distinct from other conjunctivitis with adenovirus etiology: increased ocular pressure and the presence of pseudomembranes, thin fibrin-rich films that form on the conjunctiva as a consequence of inflammatory secretions (*4-7*). HAdV-D10 has low seroprevalence across human populations making it a compelling candidate for development as a therapeutic vector (*8*).

Current therapies utilising adenovirus vectors rely heavily on species C adenovirus 5 (HAdV-C5) which has high seroprevalence across multiple populations, limiting existing therapies’ efficacy as they are neutralized by patients’ immune responses (*9*). This makes vectors based on alternative, low seroprevalence vectors more attractive as they are less likely to be recognised and neutralised by the immune system and therefore more likely to reach the target tissue (*9*). HAdV-D10 has the added benefit of low affinity receptor interaction with previously identified adenovirus receptors including Coxsackie and Adenovirus Receptor (CAR) and sialic acid, and no interaction with CD46 making it a potential ‘blank slate’ vector candidate, amenable to tissue specific targeting (*8*).

Cryo-electron microscopy (cryo-EM) has been used to determine the whole capsid structure of multiple adenoviruses including HAdV-C5, adenovirus 26 (HAdV-D26), adenovirus 41 (HAdV-F41) and the simian adenovirus ChAdOx1 derived from species E Y25 isolate, the platform vector for the ChAdOx nCoV-19 vaccine (*10-14*). Visualizing whole virus capsids during development of therapeutic vectors has previously helped with understanding off-target effects of viral vaccines and therefore may help guide engineering strategies ahead of clinical translation (*14*).

Adenoviruses have a conserved icosahedral capsid structure, consisting of three major proteins: hexon, penton base and fiber proteins. Hexon protein, the most abundant of the capsid, forms 240 trimeric capsomeres making up the facets of the icosahedron (*11, 12*). Hexon protein has seven hypervariable regions (HVRs), mostly loops, containing the most inter and intra species sequence dissimilarity (*12, 15*).

A copy of the penton base is found at each of the twelve vertices. Penton structure contains an RGD motif which is essential for engagement with host integrins to trigger virion internalisation (*16*). Penton capsomere also anchors the fiber to the capsid. The fiber is trimeric, with the shaft extending away from the capsid and terminating in the globular knob domain which acts as a high-affinity tether by interacting with host cellular receptors such as CAR, CD46 and sialic acid (SA) (*17-19*).

Minor proteins, protein (p)IIIa, pIX, pVIII and pVI, play a structural role in the capsid alongside the major proteins. Other core proteins pV, pVII and pMu (also known as pX) interact with and bind DNA in the capsid. Precursor proteins (pre-)VI, preIIIa, preTP, preVII and preMu are processed by the virally encoded protease from precursor to their mature forms (*20*). The capsid itself possesses pseudo-icosahedral symmetry as the DNA and core proteins pV, pVII and pMu do not follow strict icosahedral symmetry (*21*). Furthermore, the fiber protein located at each five-fold vertex is trimeric and has a symmetry mismatched interaction with the pentameric penton capsomere, making resolution by cryo-EM challenging. There have been few successful attempts to resolve the fiber structure of any intact adenovirus (*11*).

We pair the structure of HAdV-D10 with transcriptomic investigations to add to the body of knowledge around HAdV-D10. We use an open reading frame (ORF)-centric analysis pipeline of long read direct RNA-seq (dRNA-seq) data to cover the complex transcriptome of HAdV-D10 (*22*). This workflow has been previously used to investigate the transcriptome of adenoviruses, including HAdV-C5, ChAdOx1, an adenovirus-based vaccine against HIV and a fowl adenovirus identifying conservation of key elements of the adenovirus genome (*22-25*).

We sought to characterise the capsid structure and viral transcriptomic profile of HAdV-D10. We present the first high-resolution map of HAdV-D10 and a map of the fiber protein revealing an uncharacterized fiber shaft ‘umbrella’ motif. We observed transcription levels which varied from those observed in both HAdV-C5 and ChAdOx1 vectors, including production of expected pre-VII transcripts and additional transcripts equivalent to mature protein VII. Our study highlights the largely conserved structure of human adenovirus capsids across species while providing highly detailed transcriptomic data, adding to the portfolio of information surrounding HAdV-D10 to aid its development into a therapeutic vector.

## Results

### Structure of HAdV-D10 virions

Single particle analysis using icosahedral symmetry led to the calculation of a capsid reconstruction at 3.3 Å resolution using a total of 5,524 capsid particles. The structure of the HAdV-D10 capsid is largely similar to previously published mastadenovirus structures with global pseudo-icosahedral symmetry with a T=25 capsid. The sharply resolved map supported the assembly of models for hexon, penton, pIX, pVIII, pIIIa and pVI. To resolve the symmetry mismatched fiber and penton base, focused classification was used to first identify intact fiber shaft particles followed by focused refinement to resolve the fiber shaft at 4.6 Å resolution. Figure 1A presents the icosahedral assembly of the HAdV-D10 capsid. Figure 1B presents an exterior view of the repeating asymmetric unit of the capsid and Figure 1C presents an interior view of the asymmetric unit. The cryo-EM data used for building the atomic model are shown in Figure S1. The modelling statistics for the asymmetric unit of the HAdV-D10 capsid are listed in Table S1.

**Figure 1:**
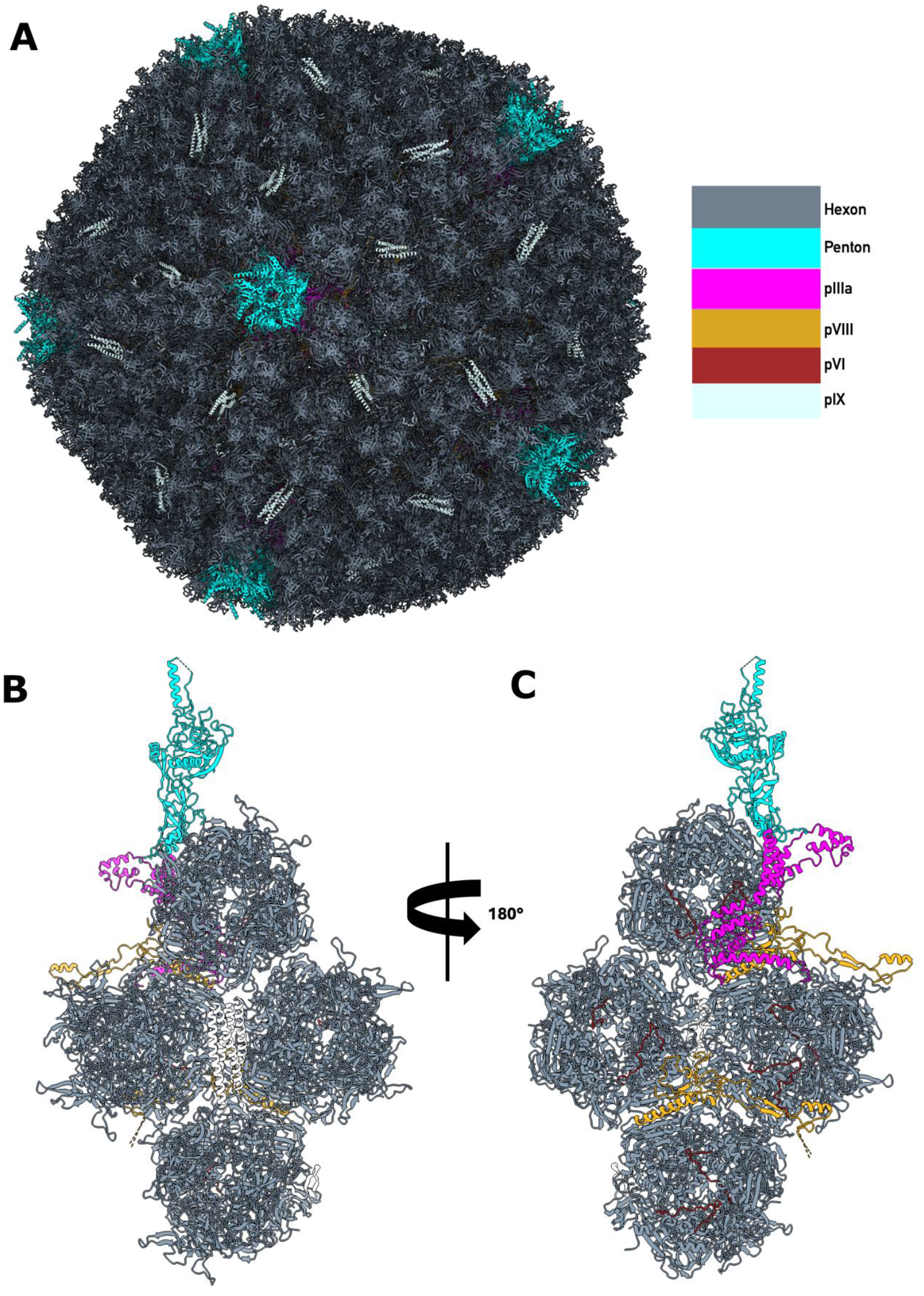
Structure Of HAdV-D10. Atomic model of biological assembly of HAdV-D10 is shown in cartoon style in panel A with a colour key highlighting protein components. Panels B and C show rotated views of the modelled asymmetric unit that repeats to form the biological assembly. Shown are exterior (B), and interior (C) capsid views respectively displaying modelled major proteins hexon and penton and minor proteins pIIIa, pVIII, pVI and pIX.

### Hexon

The full length, 949 amino acid, atomic model could be built for hexon protein. The overall architecture of hexon protein closely matches HAdV-C5 and HAdV-D26 with root mean square deviations (RSMD) of 1.158 Å and 2.448 Å respectively (supplementary Figure S3).

### Penton base

The penton base (coloured in cyan in Figure 1A-C) is highly conserved compared to HAdV-C5 (RMSD 0.762) and HAdV-D26 (RMSD 0.492) (Figure S3), and similarly is well resolved except for residues 297-324 which correspond to the loop containing the integrin binding RGD motif (Figure 1B and C).

### Fiber shaft

We used focused classification and refinement to determine the structure of the flexible fiber shaft (Figures 2 and S4). The comparatively short length of HAdV-D10 shaft at 150 Å likely reduces flexibility aiding classification. Thus, we were able to resolve the structure at 4.6 Å resolution. The length of the HAdV-D10 fiber is comparable to HAdV-D26 for which the fiber shaft structure was previously resolved (*26*). The quality of our map allowed validation of an AlphaFold3 prediction for the penton-fiber structure to be validated, fitted as a rigid body in our experimental data. We were able to experimentally demonstrate the previously uncharacterised fiber shaft motif residues 111-161. This region folds to form a shorter β-sheet, residues 133-135 and 116-119, in addition to a second longer β-sheet, residues 153-162 and 141-147. The motif has ‘confident’ Alphafold pLDDT scores of between 70-90 and matches well to the cryo-EM density (Figure 2A and B). Each β-sheet presents a solvent exposed loop that spans away from the fiber shaft to create an ‘umbrella’ like structure with the shorter β-sheet presenting a longer loop, residues 118-133, that has both electronegative and hydrophobic properties (Figure 2B-E).

**Figure 2:**
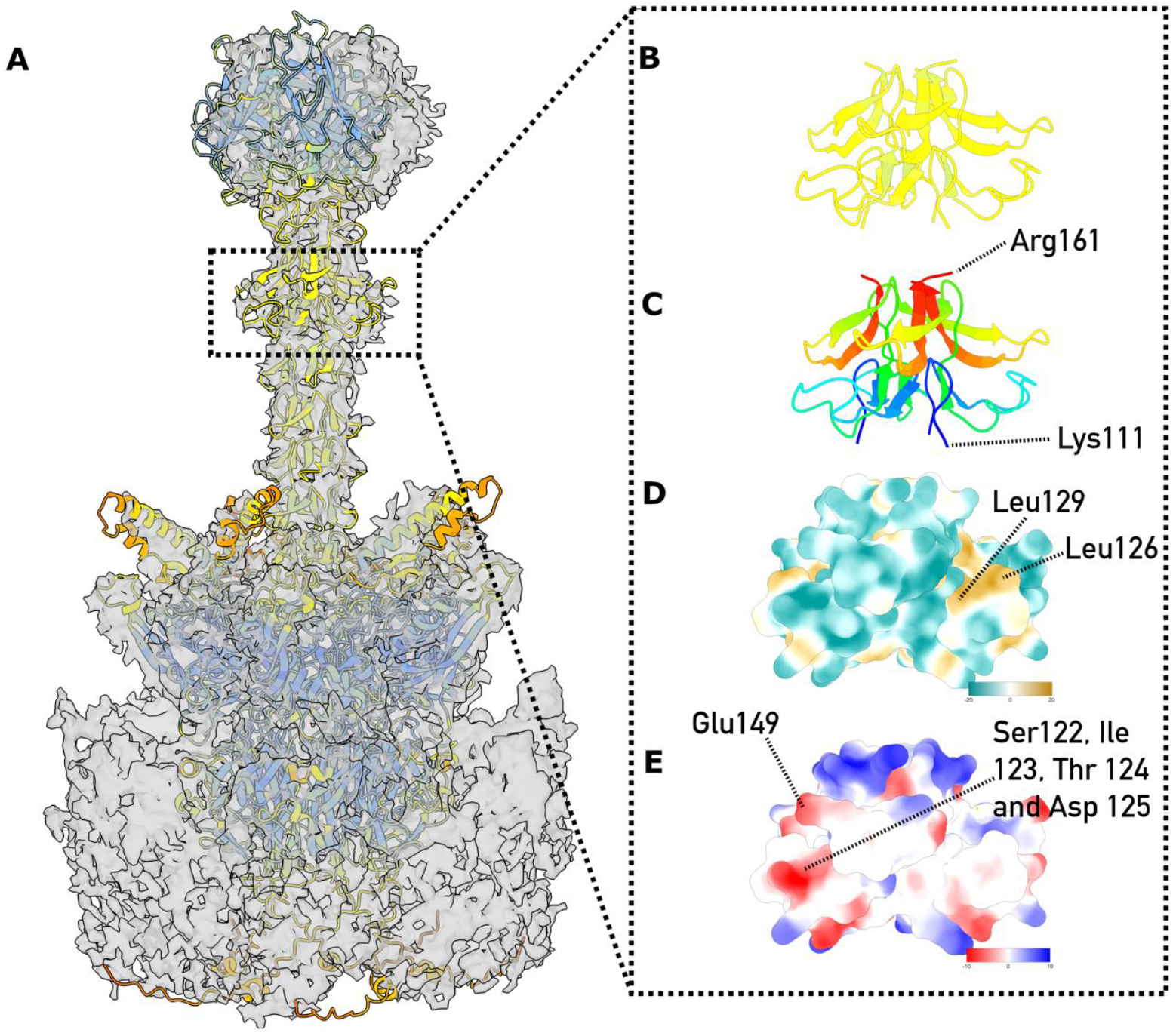
Structure Of HAdV-D10 Fiber. Panel A presents a focused refinement cryo-EM map of major protein fiber at 4.6 Å resolution with a AlphaFold3 predicted model of the trimeric fiber shaft and pentameric penton fitted into the map. The AlphaFold model is coloured by standard pLDDT confidence scores colours dark blue >90, light blue 90 - 70, yellow 70 - 50 and orange <50. Fiber ‘umbrella’ motif is highlighted with a dotted box. Panel B shows the zoomed side on view of fiber motif coloured by Alphafold pLDDT score. A view of fiber motif is presented in panel C with rainbow colour scheme to illustrate fold. N-termini to C-termini residues are coloured blue, cyan, green, yellow then red. Hydrophobic colouring and surface representation of fiber motif are presented in panel D. Coulombic colouring and surface representation of fiber motif are shown in panel E illustrating electrostatic properties of residues.

The experimental validation of this motif suggests AlphaFold3 predicted structures for other species D fiber are likely accurate. The motif is also predicted for HAdV-D26 with HAdV-D10 sharing sequence similarity of 76% when aligned structurally (Figure 3A). A similar motif is not predicted for HAdV-B35 despite the shaft being a similar length. The motif is also not predicted for HAdV-C5 (Figure 3B).

**Figure 3:**
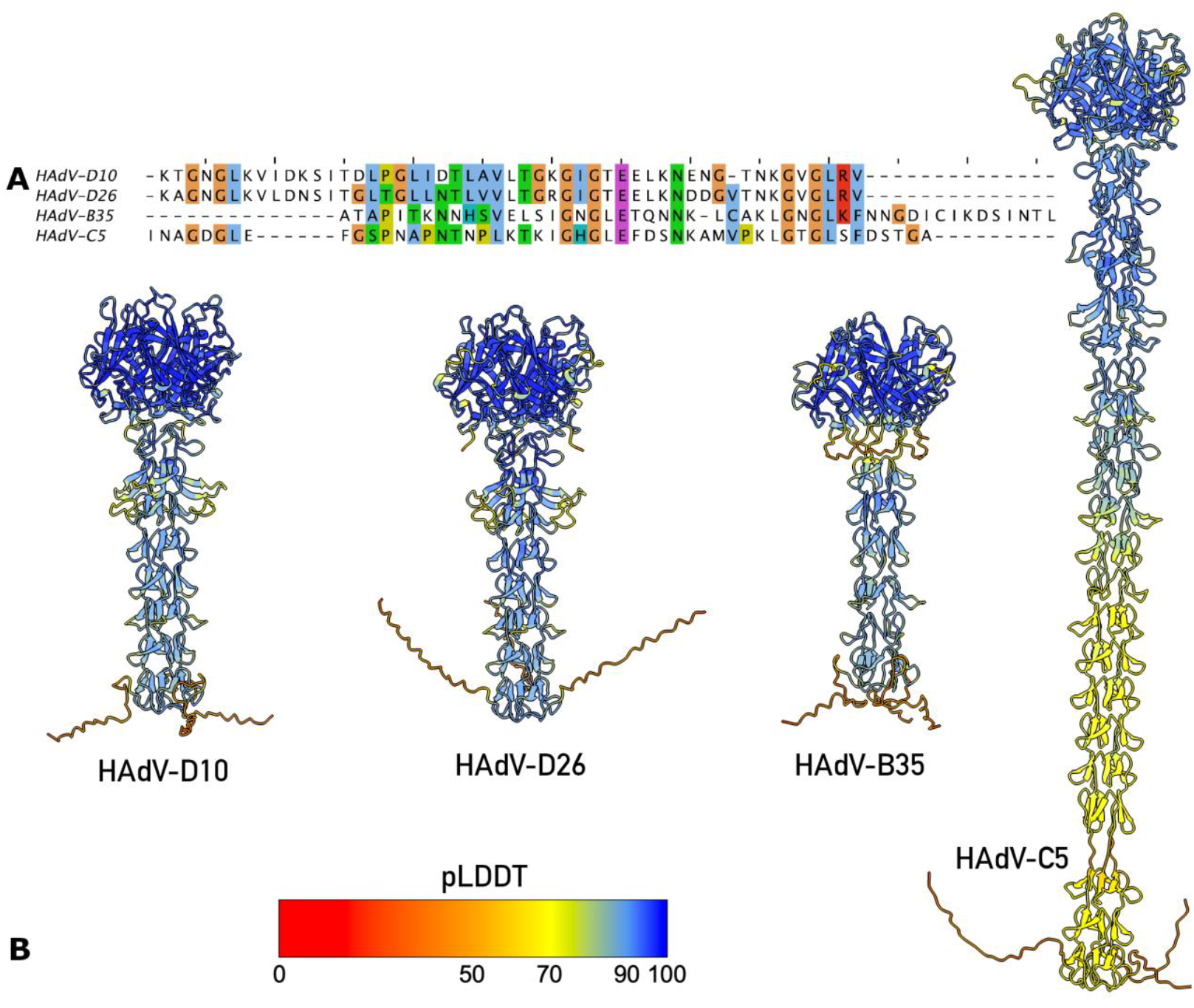
Comparison of HAdV Fiber Shaft Structures. Panel A presents sequence analysis of fiber shafts after structural alignment to umbrella motif residues 111-161 of HAdV-D10. Panel B presents a comparison of AlphaFold3 modelled structures of HAdV fiber shafts coloured by standard pLDDT confidence scores, colour key displayed. Comparison of shaft structures shows species D serotypes HAdV-D10 and HAdV-D26 have similar sequence identity and the umbrella motif predicted.

### Transcript maps of HAdV-D10 show production of key adenovirus genes

The ORF-centric analysis pipeline for the dRNA-seq data generates a transcript map showing the structure of the most abundant transcripts for each ORF present on the HAdV-D10 genome. Figure 4A presents a general overview of the classical HAdV transcriptome map with key genes highlighted and labelled with direction of transcription, aligned approximately to HAdV-D10’s genome (Figure 4B). Figure 4 presents the most abundant transcripts for each ORF for each of the 24 (Figure 4C), 48 (Figure 4D) and 72 (Figure 4E) hours post infection (h.p.i.) timepoints investigated for HAdV-D10 infected cells displayed in Integrative Genomics Viewer (IGV) (*27-29*). The maps for each of our timepoints correlates with classical adenovirus transcriptomes previously investigated, although the content of our maps differs from HAdV-C5 (*22*).

**Figure 4:**
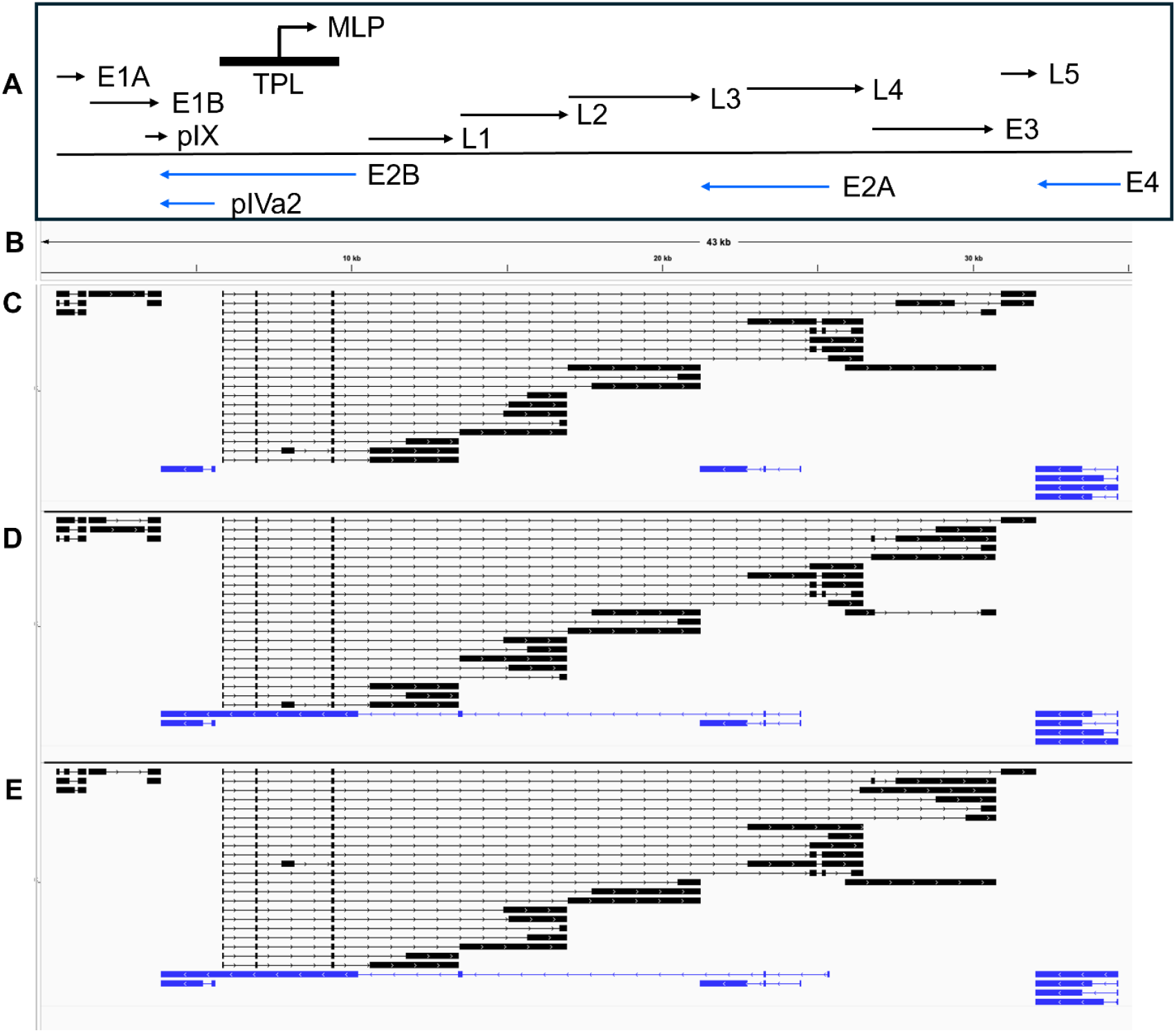
Transcription map of HAdV-D10 across multiple timepoints. Images were made in IGV (27-29). Panel A presents an overview of the classical HAdV-C5 transcriptome map with key genes highlighted (adapted from (22)). Panel B presents the HAdV-D10 genome which the following panels are aligned to. Transcript maps with the most abundant transcript for each of the known HAdV-D10 proteins are presented for 24 h.p.i. (panel C), 48 h.p.i. (panel D) and 72 h.p.i. (panel E). All transcripts are grouped according to strand, with forward strand coded transcripts shown in black and reverse strand coded transcripts shown in blue. Exons are presented as rectangles, and introns as lines, with arrows highlighting the direction of the strand.

### Change in expression of HAdV-D10 genes over time follows the expected profile

Adenovirus genes are categorized by temporal expression in the viral infection pathway, with early, intermediate and late genes being expressed sequentially (*22*). Figure 5 highlights the change in percentage of total transcripts produced per gene across the three timepoints at which RNA was sequenced. Higher percentages of early genes (E1, E2, E3 and E4 genes) are seen at 24 h.p.i. (pink bars), with increasing levels of late gene transcripts observed at 48 h.p.i. (black bars) and 72 h.p.i. (blue bars). At the end of the graph are the intermediate gene transcripts, with similar expression across the three timepoints. Supplementary Table S2 presents the percentage of each gene transcript in relation to the total number of transcripts per timepoint. This will not necessarily total 100%, due to additional transcripts arising from second potential methionine residues and transcripts that have been truncated at either the 5’ or 3’ end.

**Figure 5:**
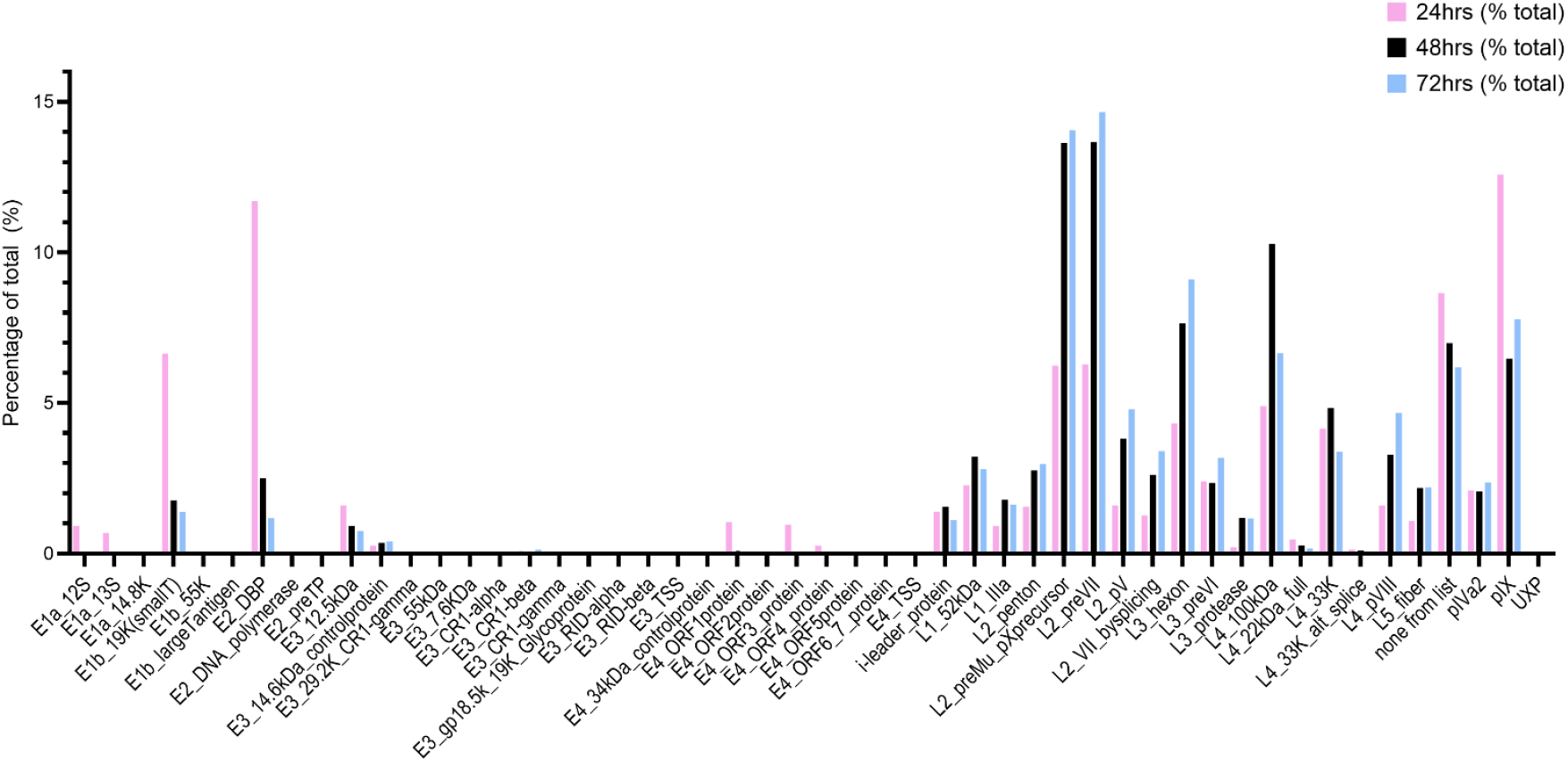
Change in transcription of known HAdV-D10 genes over 24-, 48- and 72-hours post infection. The percentage of total transcripts for each known HAdV-D10 gene are shown across the three timepoints, with values for 24 h.p.i. shown in pink, 48 h.p.i. shown in black and 72 h.p.i. shown in blue.

At the 24 h.p.i. timepoint, the gene with the highest percentage of transcripts being expressed is the intermediate gene pIX with 12.576% of total transcripts. This is closely followed by the E2 DNA binding protein with 11.703% of total transcripts at 24 h.p.i.. At the 48 and 72 h.p.i. timepoints, the L2 genes preVII and preMu are most abundant. PreVII accounts for 13.651% of all transcripts at 48 h.p.i., and 14.663% of all transcripts at 72 h.p.i. making it the most produced transcript by HAdV-D10 at later timepoints. PreMu is also highly expressed, with 13.630% of transcripts at 48 h.p.i. and 14.051% of total transcripts at 72 h.p.i..

Two E2B genes are so rarely transcribed, they account for less than 1% of total transcripts each per timepoint (*22*). The DNA polymerase was detected at each timepoint, but at such low levels that it accounts for less than 0.500% of all transcripts across the timepoints, whereas preTP has slightly increasing percentages, from less than 0.500% at 24 h.p.i., to 0.005% and 0.004% at 48 hours and 72 h.p.i..

### HAdV-D10 produces two distinct transcripts coding for precursor and mature pVII

Closer analysis of Figure 5 and Table S2 revealed two transcript groups coding for pVII were being produced. The expected preVII transcript was produced at 24, 48 and 72 h.p.i. (Figure 5), exceeding 10% of total transcripts produced in the 48 and 72 h.p.i. data sets. In contrast, a transcript coding for a protein equivalent to mature pVII was produced at less than 5% of all transcripts across all three timepoints. Figure 6 highlights the differences between these two transcript groups in the 24 h.p.i. timepoint, with the transcript coding for a protein equivalent to mature pVII being generated by a change in splice site usage (Figure 6A).

**Figure 6:**
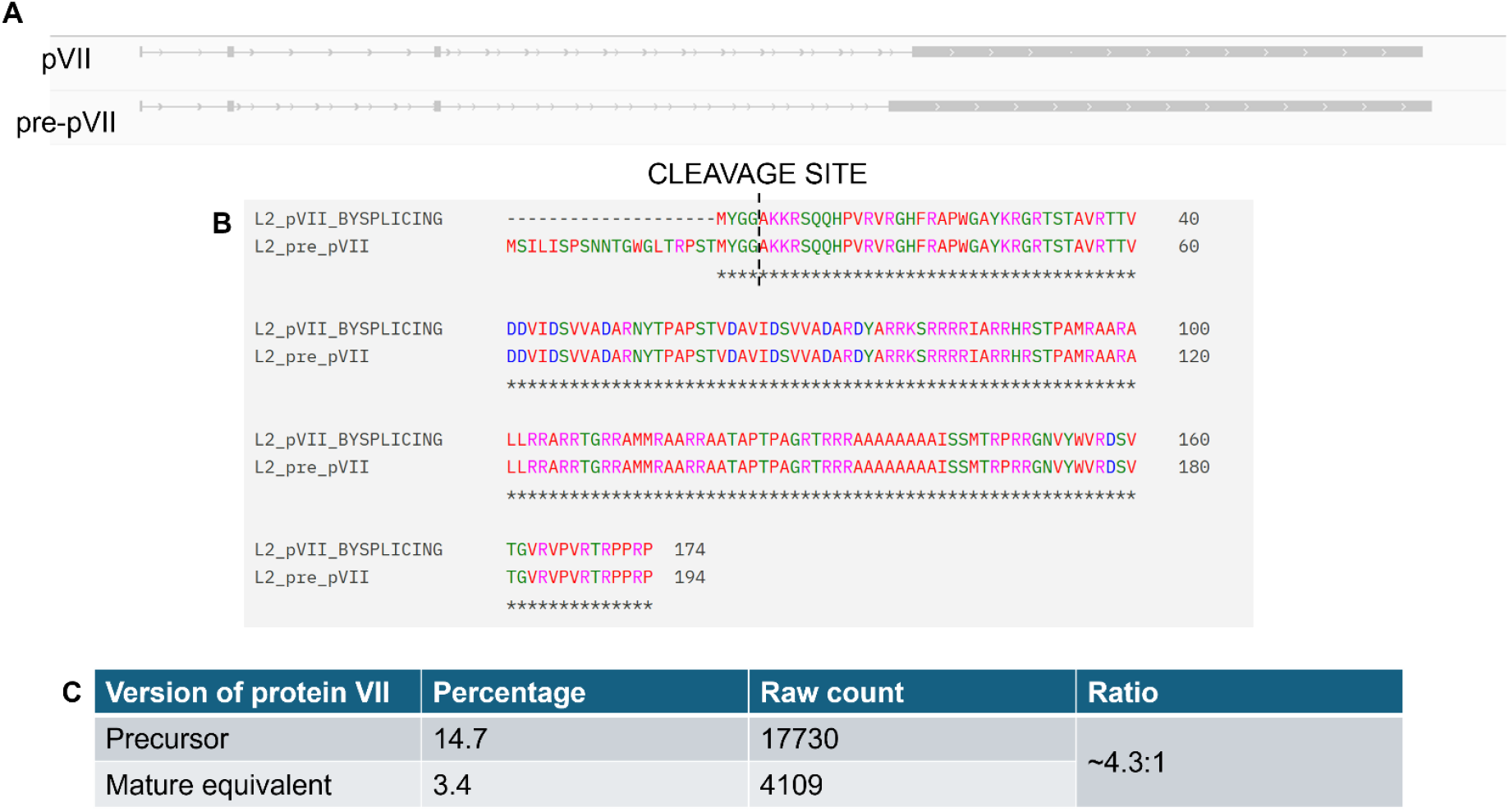
Production of preVII and pVII transcripts by HAdV-D10. Panel A illustrates HAdV-D10’s preVII and pVII transcripts observed in IGV (27-29). The transcript equivalent to mature pVII is shown first, with preVII underneath. Panel B presents the sequences of the two different transcripts, aligned by Clustal Omega (30). The cleavage site of the adenovirus protease is displayed for both of the sequences. Panel C presents the ratio of precursor to mature equivalent pVII transcripts.

The transcripts for preVII and the equivalent to mature pVII both contain the tripartite leaders, and begin at the MLP alongside other late expressed genes (Figure 4). Their sequences were aligned using Clustal Omega (*30*) and the output is shown in Figure 6B. The preVII transcript has an extension at the 5’ end, and the cleavage site for the adenovirus protease according to previous work (*31*) is highlighted after the first initiating methionine of the transcript coding for the equivalent to mature pVII, and after the second methionine for the preVII transcript.

At 72 h.p.i., preVII has a much higher raw count, at 17,730 transcripts accounting for 14.663% of all transcripts as shown in the table in Figure 6C. Although the transcript equivalent to mature pVII accounts for a smaller percentage of total transcripts, it is the 9^th^ most abundant transcript for this timepoint, exceeding the percentage for major protein coding transcripts such as the L5 fiber gene. This hints at a separate role for pVII in the HAdV-D10 lifecycle that we are currently unaware of. The two transcripts are being produced at a ratio of approximately 4.3 preVII transcripts to 1 transcript equivalent to the mature pVII.

### Hypothetical proteins from NCBI gene were identified in HAdV-D10 sequencing outputs

Figure 5 highlights a large percentage of transcripts categorized as ‘none from list’. These transcripts account for 8.600% of all transcripts at 24 h.p.i., 7.000% of all transcripts at 48 h.p.i. and 6.200% of all transcripts at 72 h.p.i.. These are large percentages compared to other transcripts being produced, with this category being the 3^rd^ most abundant transcript group at 24 h.p.i..

Using the National Centre for Biotechnology Information’s (NCBI) Blast tool (*32*), it was determined that most of these transcripts were truncated HAdV-D10 transcripts. This was not true of all the unidentified transcripts, and Table 1 presents three transcripts which were not identified as HAdV-D10 transcripts.

**Table 1:**
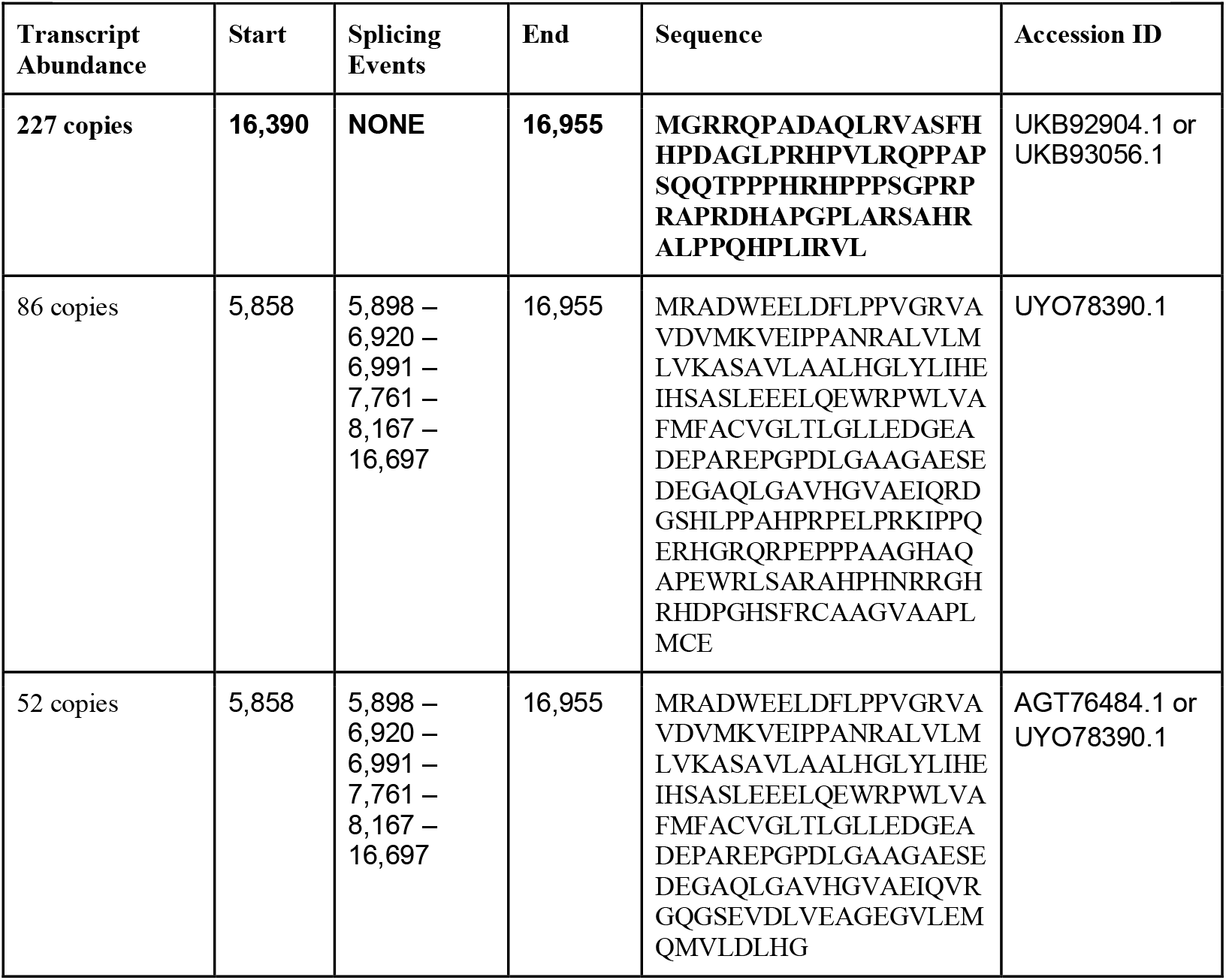
Hypothetical transcripts produced by HAdV-D10 infected cells identified by NCBI BLAST (32). Three hypothetical transcripts previously reported in NCBI Gene were identified via BLAST searching unknown transcripts. Their relative abundance has been combined for all three timepoints (labelled Transcript Abundance) and the Accession IDs associated with the hypothetical transcripts have been reported.

The start, end and splicing events are noted for each transcript, alongside their sequence and abundance across all three timepoints. After NCBI Blast searches, these were identified as hypothetical HAdV transcripts. Each Accession ID is noted in Table 1. These sequences are hypothetical, noted to have been predicted to be produced by human adenovirus genomes. The first predicted transcript listed in Table 1 is not spliced, and is initiated and terminated inside of the L2 coding region. The second and third transcripts are heavily spliced, utilising canonical GU:AG splice sites, bolstering the likelihood that they are authentic transcripts. They would be initiated at the MLP at base 5,858, and terminate in the L2 region.

## Discussion

HAdV-D10 is of interest for therapeutic applications including oncolytic virotherapies and viral vaccine vectors (*8*). Due to the infrequent nature of infections (*6*), we sought to extend the knowledge base surrounding this virus to aid further engineering and investigate its safety profile. By engineering the A20 peptide from Foot and Mouth Disease Virus (FAMDV) which binds the αVβ6 integrin (*33*) into the HAdV-D10 fiber knob protein, we were previously able to target HAdV-D10 to cells overexpressing the integrin, including several epithelial cancers (*8*). Previous literature has centered around HAdV-D10 pathology (*4-6*), therefore exploring the viral structure and transcriptomic profile enables further insights into the potential of HAdV-D10 as a therapeutic vector.

Prior structural studies have been important for identifying host interactions with viral capsid proteins (*14, 34*). In this study, we produced a 3.3 Å resolution structure of the HAdV-D10 capsid (Figures 1A, S1). This largely follows the expected structure for a human adenovirus species C and D capsids, previously determined by cryo-EM (*10-13*) as observed in Figures 1 and S3.

HAdV capsids are 80-100nm in diameter, excluding the fiber, making cryo-EM the appropriate technique for whole capsid investigation (*35*). The fiber projects away from the core capsid, anchored by the penton base as observed in Figure 2A. There are 12 fibers and pentons per capsid, found at each vertex (*11*). The fiber protein shaft is made up of variable numbers of a repeating motif, therefore the length differs across species (*36*). HAdV-D26 was observed to have a shorter, more rigid fiber shaft than HAdV-C5, making it easier to observe by cryo-EM; however the map for HAdV-D26 is not of sufficient quality to validate the predicted ‘umbrella motif’ reported here (*11*). The symmetry mismatch between the pentameric penton and trimeric fiber shaft makes resolving this anchoring interaction challenging (*37*). The 4.6 Å resolution and high quality of the fiber shaft map reported here allowed experimental validation of the AlphaFold3 structure prediction with residues 111-161 forming a motif that may act as a secondary mechanism of virus-host receptor engagement with residues Ser122, Ile123, Thr124 and Asp125 forming an electronegative patch and residues Leu126 and Leu129 forming a hydrophobic patch (Figure 2D-E). The motif presents typical features of a binding region, with a core network of hydrogen bonded residues forming a stable structure, in this case two β-sheets, that present an exposed loop to engage receptors through hydrogen bonding, electrostatic or hydrophobic interaction. For example, penton protein forms a stable hydrogen bonded α-helix, residues 282-298 reported here, that extends and projects a flexible loop into the solvent exposed space. The flexible loop contains an RGD motif that engages with host cellular integrins and corresponds to residues 299-324 in HAdV-D10 (*16*).

As further evidence serotypes should be well characterised prior to clinical translation, the transcriptomic profile of HAdV-D10 deviates from the expected profile of HAdV-C5 (*22*). Although the transcriptome maps (Figure 5C-E) align with that produced by HAdV-C5 using the same ORF-centric analysis pipeline (*22*) and an independently produced adenovirus 2 (HAdV-C2) dataset (*38*), diving into the individual transcripts and their varied production over the three timepoints revealed unexpected changes (Figure 6). While the conservation of the transcriptome map and ORFs identified in HAdV-C5 is positive from a vector development perspective, as HAdV-C5 has already been engineered as an oncolytic virus (*39*), it is important to investigate the differences as these highlight important safety considerations when developing different adenoviruses as vectors. This is highlighted by transcriptomic investigations into the ChAdOx-1 nCoV19 vaccine (*23*). Small changes from viral engineering could alter splice sites or promoter usage, introducing unintended changes. Long read sequencing confers an advantage for investigating adenovirus transcriptomics as we can investigate the high level of splicing and usage of multiple ORFs seen across adenovirus genomes, including HAdV-D10 (*40, 41*).

The differences were especially true in the case of pVII. It is normally made in a precursor form (preVII) and processed to mature pVII by the viral protease (*42, 43*). We noted some transcripts equivalent to mature pVII were produced by HAdV-D10 infected cells in addition to transcripts expected for preVII, at a ratio in the 72 h.p.i timepoint of approximately 4.3 precursor coding transcripts to 1 mature pVII coding transcript (Figure 7). Mature pVII binds DNA in the nucleoprotein core once it has been processed (*43, 44*). HAdV-D10 infected cells are producing a far higher percentage of total reads for preVII and pVII, compared with major structural protein coding transcripts such as the genes encoding the hexon and fiber (Figures 5&6), suggesting that, as in HAdV-C5, the stoichiometry of gene transcripts to genes does not reflect the number of protein copies in a viral capsid (*22*).

We revisited the HAdV-C5 dataset and a single transcript coding for mature pVII was found (*22*), which may suggest an undetermined role or increased need for pVII by HAdV-D10 to explain the increased number of transcripts produced. pVII has been described as a histone-like protein and is thought to have roles including evading the host immune system, and modulating the host’s DNA damage response during HAdV infection (*45, 46*). Figure 5 also highlighted a higher percentage of total transcripts for the gene encoding preMu, which has been suggested to modulate splice site selection during adenovirus infection (*47*). This protein is also involved in binding DNA and localises in the nucleoprotein core, as with preVII/pVII (*44, 47*) strengthening the position that HAdV-D10 may express DNA binding genes at higher levels relative to HAdV-C5. DNA condensation is extreme in adenoviral capsids. Large amounts of DNA must be packed into the 80-100nm icosahedral capsid and pMu, pVII and pV have been reported to work similarly to eukaryotic histone proteins to condense adenovirus DNA, interacting with each other to form adenosomes which can be observed via cryo-electron tomography (cryo-ET) (*21, 42*).

Why HAdV-D10 might require relatively higher levels of DNA binding proteins compared to HAdV-C5 requires more exploration, to elucidate the interplay of the different proteins produced by HAdV-D10 in host cells. Proteomic investigations may aid this, to confirm if the high number of transcripts correlates with levels of protein production, or if the stoichiometry is unequal as seen in HAdV-C5 (*22*). Proteomic analysis would also help confirm production of the hypothetical transcripts identified in Table 1 are protein encoding, as performed previously alongside dRNA-seq data (*40*).

In conclusion, we have combined the first whole capsid structure of the HAdV-D10 capsid at 3.3 Å (Figures 1 and S1) with the first transcriptomic investigations (Figure 4) of the same virus, revealing unique features and insights into its biology. We have also provided a 4.6 Å map of the fiber shaft revealing a previously uncharacterised motif (Figure 2). While mostly sharing structural characteristics to HAdV-C5 (*10*), there are key variations in the transcriptomic data including the presence of hypothetical transcripts and unexpected transcripts equivalent to mature pVII (Figure 6 and Table 1). While these differences require further proteomic investigation, HAdV-D10 remains a promising potential vector for therapeutic application. The combined structural and biological information presented in this work can be used to improve safety and efficacy of future HAdV-D10-based vectors and guide vector engineering strategies.

## Materials and Methods

### Virus production and purification

All viruses were produced as previously described (*48*). Briefly, viruses were produced in E1A complementing cell lines and were purified by multiple rounds of caesium chloride (CsCl) gradient ultracentrifugation before dialysis against a buffer made up of 10% glycerol, 135mM NaCl, 10mM Tris-HCl (pH 7.8) and 1mM MgCl_2_(H_2_O)_6_. A viral titre was determined by microBCA assay (Pierce microBCA protein assay kit, ThermoFisher Scientific, Waltham, USA, Catalogue number: 23235) and by immunostaining assay to determine the functional titre. To convert the microBCA result to a viral titre, the formula 1µg protein = 4 × 10^9^ viral particles/mL was used (*49, 50*). The wild-type HAdV-D10 virus used for transcriptomic experiments had a measured functional titre of 3.26×10^10^ PFU/mL while the replication deficient HAdV-D10 virus used for cryo-EM had a viral titre of 3.02×10^12^ VP/ml by microBCA assay.

### Cryo-EM grid preparation

Grids of purified HAdV-D10 were prepared using Quantifoil R2/2 Cu 300 mesh grids, covered in-house with a thin layer of carbon produced using a Quorum Q150. Grids were glow discharged using a Emitech K100K for 30 seconds at 15 mA in air. Plunge freezing was performed using a Vitrobot Mark IV into liquid ethane. The Vitrobot chamber was set to 95% humidity and 4.5°C. 6 ul of sample was applied and, after a 60 second wait time, blotted for 6 seconds with a blot force of −6.

### Cryo-EM data collection

Data collection for the high-resolution reconstruction was performed at Diamond Light Source Electron Bio-Imaging Centre (eBIC) using a Thermofisher Titan Krios equipped with a Gatan K3 detector operated in super resolution mode. Exposures with a total dose of 40 e/Å2 were recorded over 40 frames, and a nominal defocus range of −2.2 µm to −0.2 µm. In total 22,490 micrographs were collected. The recorded images were binned by two, resulting in a calibrated pixel size of 0.829 Å/pixel at the image level.

### Single particle data analysis of HAdV-D10 Capsid

Single particle analysis was performed in RELION 3.1 using a high-performance computing cluster (HPC) with GPU acceleration from NVIDIA P100’s. Briefly, motion correction was performed using MotionCorr2. Contrast transfer function was estimated using GCTF, with poorly estimated micrographs, based on the fit of the real CTF to the theoretical CTF, removed from further processing by visual inspection of the Thon rings.714 micrographs were removed at this stage. 100 particles were manually picked and used as a 2D class templates for RELION’s “Autopicker” resulting in a dataset set of 11,344 particles. Particle false positives were removed by multiple iterative rounds of 2D classification. An ab-initio model was generated from the data using the RELION’s stochastic gradient descent algorithm without the imposition of symmetry. The Ab-initio model was subsequently aligned to I2 symmetry using “relion_align_symmetry.” The I2 aligned Ab-initio reference map was used as a reference to refine particles at 5x, 4x and subsequently 3x Fourier cropping factors, re-extracting the particles after each refinement reached the Nyquist limit of the Fourier cropped pixel size. Particles were then re-extracted in a box of 1440 pixels (∼1194 Å) Fourier cropped by a factor of two to 720 pixels. This was the maximum box size allowed within memory constraints of the HPC. After multiple rounds of iterative CTF and 3D refinement with per-particle defocus, astigmatism, beamtilt and trefoil the resulting map has a resolution of 3.8 Å with 5,524 particles included in the reconstruction. After Ewald sphere correction, the resolution estimate for the capsid by gold standard Fourier shell correlation (FSC) calculations is 3.3 Å.

### Model building

“Phenix_sharpen” was used to sharpen the unfiltered half maps (*51*). SwissModel (*52*) and the Alphafold3 server (*53*) were used to generate homology models for the capsid proteins. Each homology model was first fitted into the map using ChimeraX’s “fit in map” function as rigid bodies (*54*). The models were then independently adjusted, particularly for C-α backbone fit into data, using COOT (*55*). Once all models with experimental data were independently built, ISOLDE was used to refine the whole asymmetric unit, first allowing the asymmetric unit model to relax into the map by running an all-chain molecular dynamics AMBER forcefield simulation with a temperature of 20 K. Subsequently the temperature was lowered from 20 K to 1 K and individual residues were inspected to improve side chain fit to data, rotamer angles and Cα Phi and Psi angles (*56*). Finally, one round of phenix real space refinement was performed to generate the.cif file. The biological icosahedral assembly, shown in Figure 1A, was produced by symmetry-related copies using Chimera’s “Sym” command.

### Focused classification and refinement of fiber shaft

The fiber shaft map was determined using 3D classification and refinement without the application of symmetry (C1) and targeted masking within RELION 3.1. First, the refined particles from the 3.3 Å global capsid map were re-extracted and Fourier cropped by a factor of 5 from the previously used box size of 1440 pixels (1194 Å). “Refine3D” was then used to output a particle.star file which was symmetry expanded by icosahedral symmetry using “relion_symmetry_expand.” A cylindrical mask that covered the diameter of the penton and spanned the expected length of the HAdV-D10 fiber shaft was made using MATLAB and DYNAMO (*57*). The mask was resampled onto the grid of the reference map. RELION “MaskCreate” was then used to apply a soft edge to the cylindrical mask. “3D Classification” was then performed using the expanded symmetry particles, Fourier cropped reference map and cylindrical mask without performing image alignment. The resulting classes were assessed for intact fiber shaft particles and re-extracted in a 1440 pixel box Fourier cropped to 720 pixels. Using the intact fiber shaft particle coordinates “Particle Subtraction” was performed using the same cylindrical mask in a new smaller box size of 200 Å. 3D classification was performed with image alignment, using a regularisation parameter of 4, performing local angular searches from 5 degrees, for 25 iterations and filtering particles into 5 classes. 3 classes had full length fiber shaft particles and were selected for further refinement. The subsequent converged refinement was sharpened by b-factor using RELION’s “PostProcess” resulting in the final map shown in Figure 2A and S4.

### AlphaFold structure prediction

All AlphaFold predictions reported here were generated using the AlphaFold3 server (*53*). For the Fiber-Penton model, five copies of the penton and three copies of the fiber sequence were submitted to the AlphaFold3 server as two separate protein entities. The model was then manually placed into the cryo-EM map and then fitted using ChimeraX’s “fit in map” command.

### RNA extraction, mRNA enrichment and sequencing

Kyse-30 cells were grown to a minimum of 80% confluency in T75 flasks. Cells were infected with wild-type HAdV-D10 at an MOI (multiplicity of infection) of 10 determined by immunostaining assay.

RNA was extracted 24 h.p.i. using previously described methods utilising TRI reagent (ThermoFisher Scientific, Waltham, USA, Catalogue number: AM9738) and chloroform (*22*). The resulting pellet following isopropanol precipitation was washed three times with ethanol to increase purity.

The DynaBeads mRNA Purification kit (ThermoFisher Scientific, Waltham, USA, Catalogue number 61006) was used to prepare total RNA extracted for polyA-tail enrichment according to manufacturers’ instructions. Resulting mRNA was then used for sequencing. The dRNA-seq kit from Oxford Nanopore Technologies (Oxford, UK, Catalogue number: SQK-RNA004) was used according to manufacturers’ instructions to prepared dNRA-seq libraries and perform sequencing using a MinION Mk1C (Oxford Nanopore Technologies, Oxford, UK, Catalogue number M1CBASICSP) and RNA flow cells (Oxford Nanopore Technologies, Oxford, UK, Catalogue number: FLO-MIN004RA). Data was acquired over 72 hours per sample achieving the following number of reads for each run: 876.140 thousand for 24 h.p.i., 2.290 million for 48 h.p.i. and 3.040 million for 72 h.p.i..

### Transcriptomics data analysis

An ORF (open reading frame)-centric pipeline was applied to the data output from the MinION Mk1C as previously described (*22*). Reads were mapped to the HAdV-D10 genome using minimap2 (*58*) to filter out host and positive control transcripts.

The mapped transcripts were then counted and classified, using a previously described script: “classify_transcripts_and_polya_segmented_V2.pl” (*22*). These groups were then named using a second script: “name_transcripts_and_track.pl” (*22*) which utilises a table of features expected to be found in the HAdV-D10 genome and their expected transcription start site to produce a list of transcripts and which proteins they could produce if they were translated. Outputs were visualised in IGV (*27-29*) and graphs produced in GraphPad Prism version 10.4.1 (627) (*59*). All scripts used for transcriptomic data analysis are available to access from Zenodo (*60*).

### Alignment of pVII transcripts

Transcript sequences for pVII on both the mature equivalent and precursor form were obtained from the dRNA-seq analysis output. They were aligned using Clustal Omega (*30*).

## Supporting information

Mundy, Waraich et al. Supplemental Files

## Acknowledgments

We acknowledge Diamond for access and support of the cryo-EM facilities at the UK national electron bio-imaging centre (eBIC), proposal BI31827-1. The authors would like to thank Lorna Malone for assistance as the local contact at eBIC. We would also like to thank Mairi Clarke, Kako Stapleton and James Streetley from the Scottish Centre of Macromolecular Imaging (SCMI) for help with preliminary cryo-EM data.

## Funding

R.M.M. was supported in part by grant MR/N0137941/1 for the GW4 BIOMED MRC DTP, awarded to the Universities of Bath, Bristol, Cardiff and Exeter from the Medical Research Council (MRC) and in part by Advanced Therapies Wales. KW was supported by a JEOL-University of Glasgow Industrial Partnership Studentship awarded to D.B. EAB was funded by a Cardiff University Ph.D. studentship to A.L.P., by the Experimental Cancer Medicine Centre award to Cardiff University (reference C7838/A25173) and this work was supported by The Brain Tumour Charity (grant number GN-000755). We acknowledge the Scottish Centre for Macromolecular Imaging (SCMI) for access to cryo-EM instrumentation, funded by the MRC (MC_PC_17135, MC_UU_00034/7, MR/X011879/1) and SFC (H17007).

## Author contributions

RMM – Investigation, Methodology, Formal analysis, Data curation, Visualisation, Writing – original draft, Writing review & editing; KW - Investigation, Methodology, Formal analysis, Data curation, Visualisation, Writing – original draft, Writing review & editing; EAB – Investigation, Methodology, Resources, Writing review & editing; PJR – Validation, Formal analysis, Investigation, Supervision, Writing review & editing; ATB – Methodology, Formal analysis, Writing review & editing; MTY – Formal analysis, Writing review & editing; EM – Methodology, Investigation, Formal analysis, Supervision, Writing review & editing; PdF – Methodology, Investigation, Formal analysis, Supervision, Writing review & editing; CMB – Formal analysis, Resources, Supervision, Writing review & editing; DM – Conceptualisation, Methodology, Software, Validation, Formal analysis, Resources, Supervision, Writing review & editing; DB - Conceptualisation, Methodology, Validation, Formal analysis, Resources, Supervision, Writing review & editing; ALP - Conceptualisation, Methodology, Validation, Formal analysis, Resources, Supervision, Writing – original draft, Writing review & editing, Funding acquisition.

## Competing interests

ALP is CSO of Trocept Therapeutics (part of Accession Therapeutics Ltd) who are developing adenoviral platforms for clinical oncology applications. Accession Therapeutics Ltd had no role in this manuscript. All other authors declare no conflict of interest.

## Data and materials availability

The HAdV-D10 capsid model is deposited in the Protein Data Bank accession code 9R78. The associated capsid map is deposited in the Electron Microscopy Databank (EMDB) accession code EMDB-53736. The fiber shaft map is deposited in the EMDB accession code EMDB-53655.

The scripts used for analysis of transcriptomic data are available from Zenodo (*60*) (https://doi.org/10.5281/zenodo.3610257).

