## Supplementary material for "Identification of a novel structural motif and overexpression of key transcripts elucidated in Adenovirus 10": Mundy, Waraich et al. Supplemental Files

**This PDF file includes:**

Figs. S1 – S4
Tables S1 to S3

**Fig. S1.**

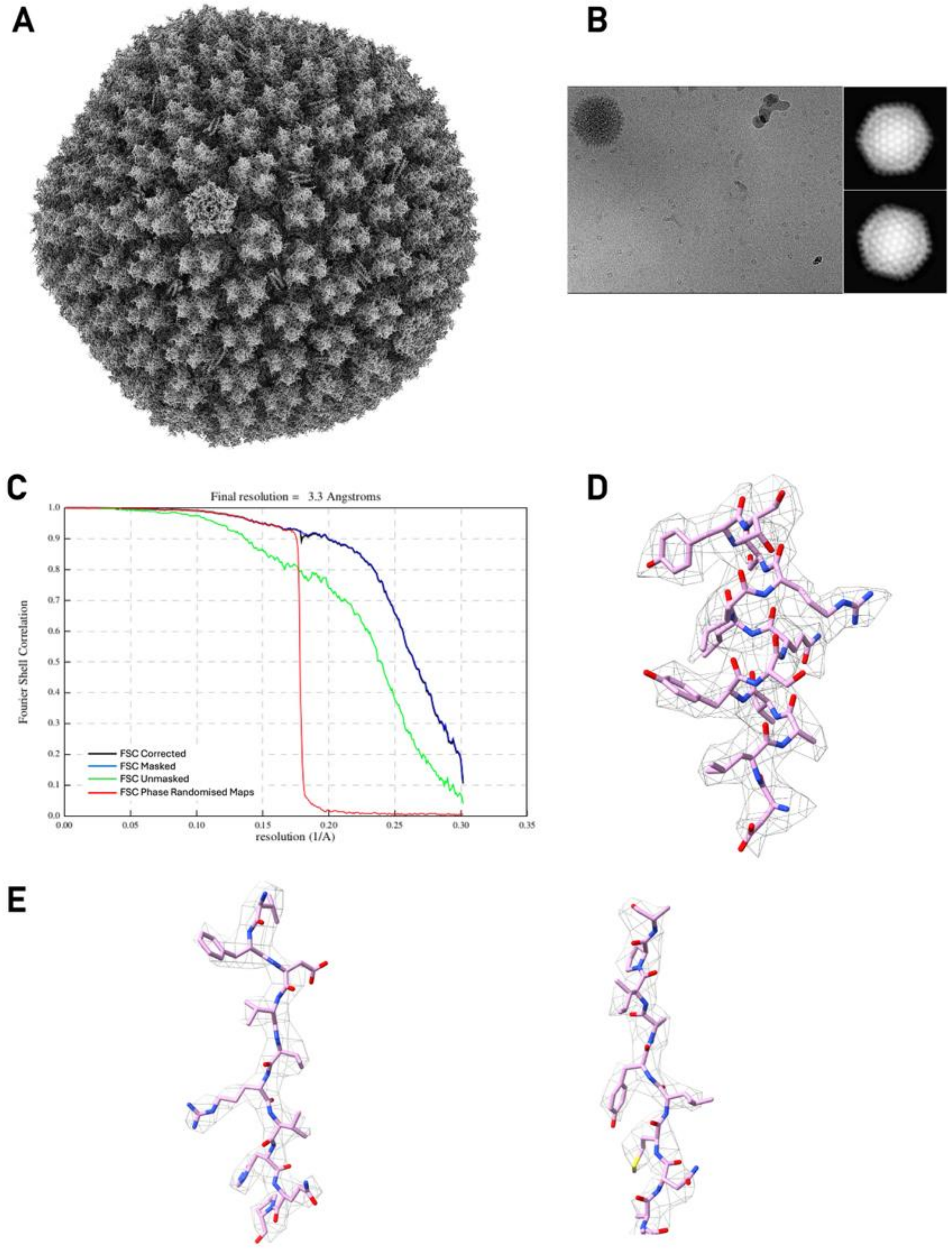


**Figure S1**: **Cryo-EM data of HAdV-D10 capsid. A** Cryo-EM map of HAdV-D10 capsid at 3.3 Å resolution. **B** Example micrograph and 2D classes. In total 22,490 micrographs were collected at a nominal magnification of 105k. On average there is ~1 particle per micrograph with at least 11,344 virion particles in the data collection. **C** Fourier Shell Correlation (FSC) plots for Ewald sphere corrected map. **D** Example of model fitting to cryo-EM data with atomic model chain displayed in stick representation. Residues shown are penton 426-438. **E** Example of hexon model fitting to cryo-EM data, residues 646-655 and 919-928 are shown.

**Fig S2.**

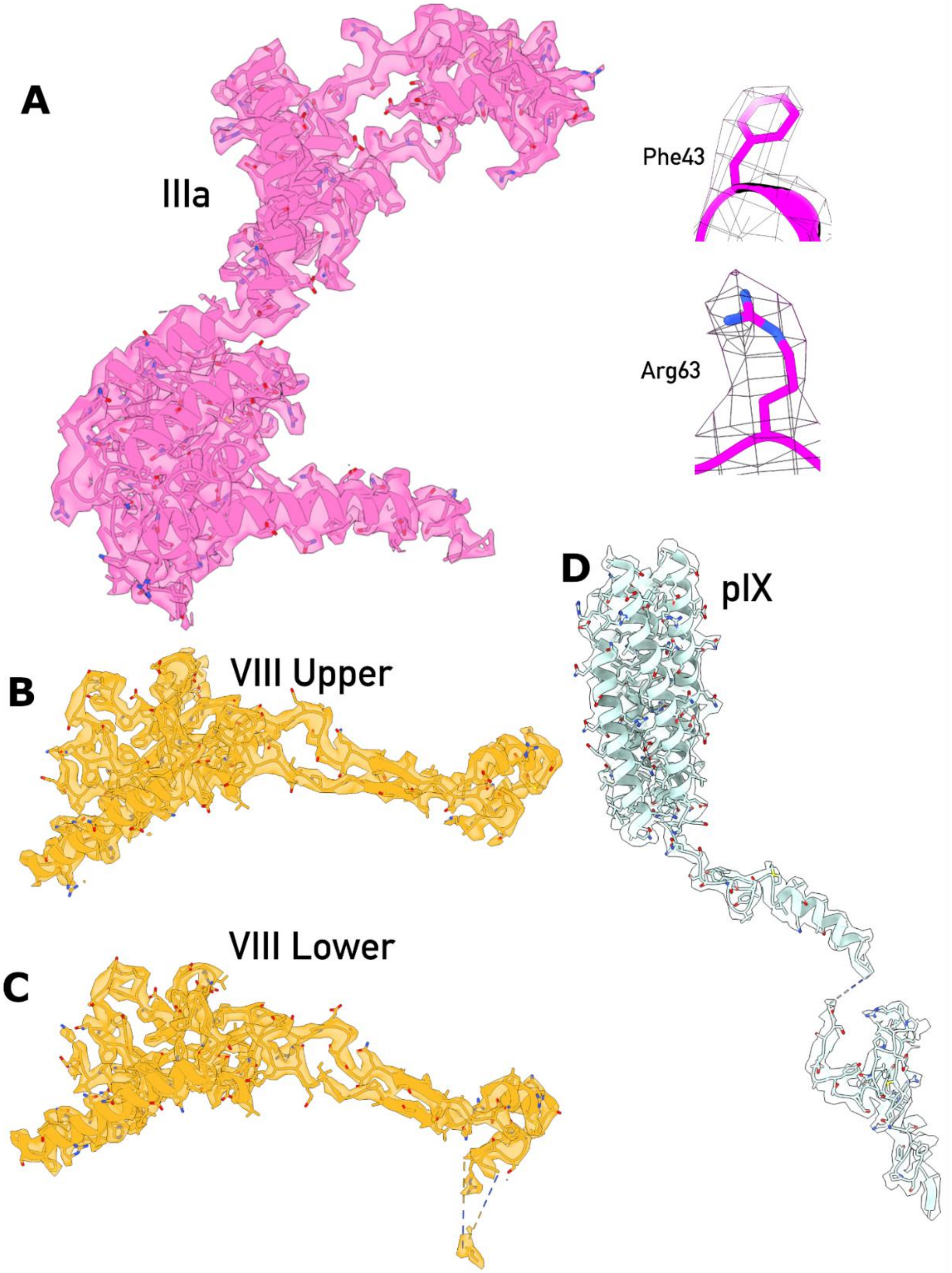


**Figure S2. Structure Of Capsid Minor Proteins**. Despite the low particle number included in the final reconstruction the quality of the map density for the minor proteins is strong with many side chains modelled accurately particularly for pIIIa (panel A), evidence the HAdV-D10 is a more stable capsid than previously studied adenovirus serotypes. Panels B and C present upper and lower pVIII structures. pIX (panel D) forms a coiled-coil domain as observed in HAdV-C5 and HAdV-D26.

**Fig. S3.**

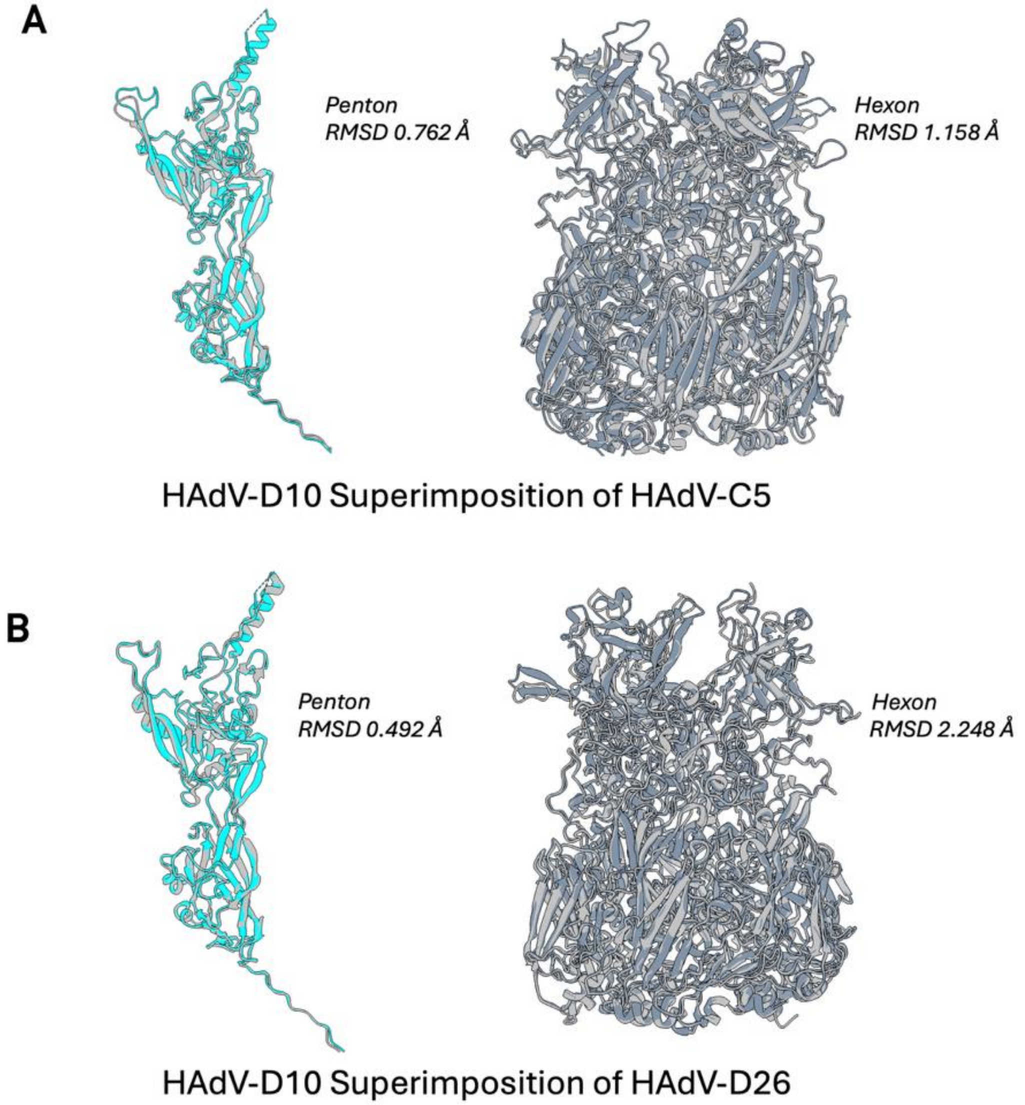


**Figure S3: Superimposition of HAdV-D10 for comparison to HAdV-C5 and HAdV-D26.** ChimeraX's MatchMaker command was used to first perform structural alignment and then calculate least-squares-fit root mean-square-deviations (RMSD) of Cα backbone residues. Colour scheme from Figure 1 for HAdV-D10 penton and hexon is maintained. **A** Comparison of HAdV-D10 penton, (cyan), and hexon (slate grey) with HAdV-C5. **B** Comparison of HAdV-D10 penton (cyan) and hexon (slate grey) with HAdV-D26.

**Fig. S4.**

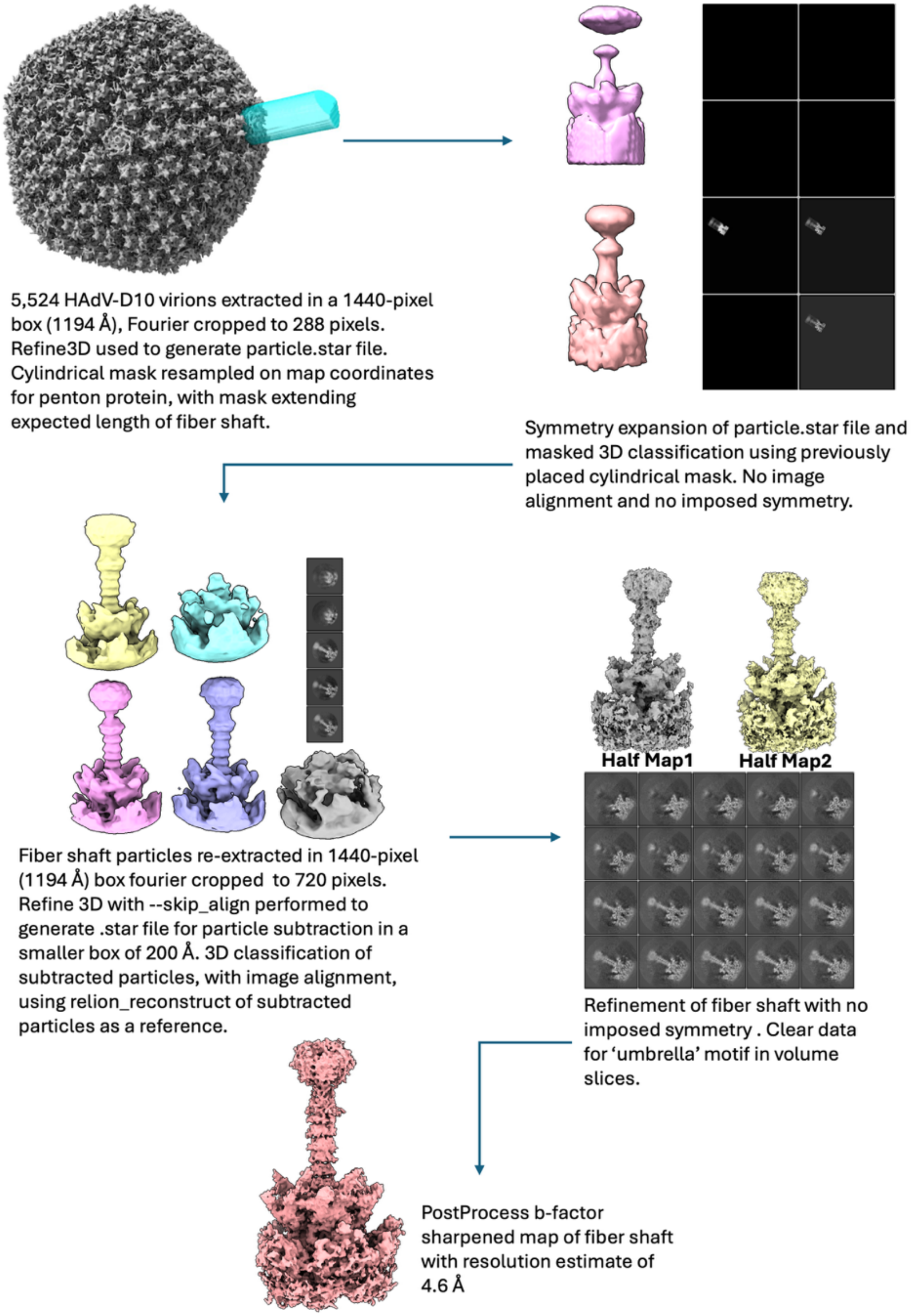


**Figure S4. Processing pipeline for focussed refinement of fiber shaft.**

**Table S1: Modelling statistics for HAdV-D10 capsid asymmetric unit.**

| | |
|---|---|
| Clashscore | 1.18 |
| Poor rotamers | 171 / 1.56% |
| Favoured rotamers | 10399 / 94.58% |
| Ramachandran outliers | 99 / 0.78% |
| Ramachandran favoured | 11677 / 92.14% |
| Rama distribution Z-score | -1.88 ± 0.07 |
| MolProbidity score | 1.46 |
| Cβ deviations >0.25 A | 101 / 0.85% |
| Bad Bonds | 0 of 103946 / 0.00% |
| Bad Angles | 824 of 141412 / 0.58% |
| Cis Prolines | 24 of 816 / 0.05% |
| Cis nonProlines | 6 of 11890 / 0.02% |
| Twisted Peptides | 2 of 12706 / 0.02% |
| CaBLAM outliers | 486 / 3.9% |
| CA Geometry outliers | 203 / 1.61% |

**Table S2: Percentage of each transcript produced across 24, 48 and 72 h.p.i. by cells infected with HAdV-D10.**

| Feature | Percent of total (24hr) | Percent of total (48hr) | Percent of total (72hr) |
|---|---|---|---|
| E1a_12S | 0.917 | 0.0516 | 0.0521 |
| E1a_13S | 0.6986 | 0.0478 | 0.0678 |
| E1a_14.8K | 0.0436 | 0.0151 | 0.0157 |
| E1b_19K(smallT) | 6.6375 | 1.7743 | 1.3861 |
| E1b_55K | 0 | 0.0012 | 0 |
| E1b_largeTantigen | 0 | 0 | 0 |
| E2_DBP | 11.703 | 2.5064 | 1.1851 |
| E2_DNA_polymerase | 0 | 0 | 0 |
| E2_preTP | 0 | 0.005 | 0.0041 |
| E3_12.5kDa | 1.6157 | 0.9161 | 0.7542 |
| E3_14.6kDa_controlprotein | 0.262 | 0.3616 | 0.406 |
| E3_29.2K_CR1-gamma | 0 | 0.0315 | 0.0306 |
| E3_55kDa | 0 | 0 | 0 |
| E3_7.6KDa | 0 | 0 | 0 |
| E3_CR1-alpha | 0 | 0 | 0.0008 |
| E3_CR1-beta | 0.0873 | 0.0831 | 0.1273 |
| E3_CR1-gamma | 0 | 0 | 0 |
| E3_gp18.5k_19K_Glycoprotein | 0 | 0.0025 | 0 |

| | | | |
|---|---|---|---|
| E3_RID-alpha | 0 | 0 | 0 |
| E3_RID-beta | 0 | 0 | 0.0016 |
| E3_TSS | 0 | 0 | 0 |
| E4_34kDa_controlprotein | 0 | 0 | 0 |
| E4_ORF1protein | 1.048 | 0.0919 | 0.0719 |
| E4_ORF2protein | 0.0436 | 0.0201 | 0.0124 |
| E4_ORF3_protein | 0.9606 | 0.0655 | 0.0504 |
| E4_ORF4_protein | 0.262 | 0.0466 | 0.0396 |
| E4_ORF5protein | 0 | 0 | 0 |
| E4_ORF6_7_protein | 0 | 0 | 0 |
| E4_TSS | 0 | 0 | 0 |
| i-leader_protein | 1.3973 | 1.5575 | 1.1239 |
| L1_52kDa | 2.2707 | 3.2234 | 2.8003 |
| L1_IIIa | 0.917 | 1.7869 | 1.6375 |
| L2_penton | 1.572 | 2.7622 | 2.974 |
| L2_preMu_pXprecursor | 6.2445 | 13.6298 | 14.0505 |
| L2_preVII | 6.2882 | 13.6513 | 14.6634 |
| L2_pV | 1.6157 | 3.8119 | 4.7902 |
| L2_VII_bysplicing | 1.2663 | 2.6186 | 3.3983 |
| L3_hexon | 4.3231 | 7.6453 | 9.1065 |
| L3_preVI | 2.4017 | 2.3464 | 3.1816 |
| L3_protease | 0.2183 | 1.1971 | 1.171 |

| | | | |
|---|---|---|---|
| L4_100kDa | 4.8908 | 10.2728 | 6.6585 |
| L4_22kDa_full | 0.4803 | 0.2847 | 0.1695 |
| L4_33K | 4.1484 | 4.8364 | 3.3784 |
| L4_33K_alt_splice | 0.131 | 0.0995 | 0.0686 |
| L4_pVIII | 1.6157 | 3.2902 | 4.6769 |
| L5_fiber | 1.0917 | 2.1838 | 2.2024 |
| none from list | 8.6462 | 6.9913 | 6.1829 |
| pIVa2 | 2.096 | 2.0654 | 2.3636 |
| pIX | 12.5764 | 6.4721 | 7.7832 |
| UXP | 0 | 0 | 0 |

**Table S3: Hypothetical transcripts produced by HAdV-D10 infected cells identified by NCBI BLAST (*32*).** Three hypothetical transcripts previously reported in NCBI Gene were identified via BLAST searching unknown transcripts. Their relative abundance has been combined for all three timepoints (labelled Transcript Abundance) and the Accession IDs associated with the hypothetical transcripts have been reported.

| Transcript Abundance | Start | Splicing Events | End | Sequence | Accession ID |
|---|---|---|---|---|---|
| 227 copies | 16,390 | NONE | 16,955 | MGRRQPADAQLRVASFHHPDAGLPRHPVLRQPPAPSQQTPPPHRHPPPSGPRPRAPRDHAPGPLARSAHRALPPQHPLIRVL | UKB92904.1 or UKB93056.1 |
| 86 copies | 5,858 | 5,898 – 6,920 – 6,991 – 7,761 – 8,167 – 16,697 | 16,955 | MRADWEELDFLPPVGRVAVDVMKVEIPPANRALVLMLVKASAVLAALHGLYLIHEIHSASLEEELQEWRPWLVAFMFACVGLTLGLLEDGEADEPAREPGPDLGAAGAESEDEGAQLGAVHGVAEIQRDGSHLPPAHPRPELPRKIPPQERHGRQRPEPPPAAGHAQAPEWRLSARAHPHNRRGHRHDPGHSFRCAAGVAAPLMCE | UYO78390.1 |
| 52 copies | 5,858 | 5,898 – 6,920 – 6,991 – 7,761 – 8,167 – 16,697 | 16,955 | MRADWEELDFLPPVGRVAVDVMKVEIPPANRALVLMLVKASAVLAALHGLYLIHEIHSASLEEELQEWRPWLVAFMFACVGLTLGLLEDGEADEPAREPGPDLGAAGAESEDEGAQLGAVHGVAEIQVRGQGSEVDLVEAGEGVLEMQMVLDLHG | AGT76484.1 or UYO78390.1 |
